# Region-specific and dose-specific effects of chronic haloperidol exposure on [^3^H]-Flumazenil and [^3^H]-Ro15-4513 GABA_A_ receptor binding sites in the rat brain

**DOI:** 10.1101/869941

**Authors:** Alba Peris-Yague, Amanda Kiemes, Diana Cash, Marie-Caroline Cotel, Nisha Singh, Anthony C. Vernon, Gemma Modinos

## Abstract

*Post-mortem* studies suggest that schizophrenia is associated with abnormal expression of specific GABA_A_ receptor (GABA_A_R) α subunits, including α5GABA_A_R. Positron emission tomography (PET) measures of GABA_A_R availability in schizophrenia, however, have not revealed consistent alterations *in vivo*. Animal studies using the GABA_A_R agonist [^3^H]-muscimol provide evidence that antipsychotic drugs influence GABA_A_R availability, in a region-specific manner, suggesting a potential confounding effect of these drugs. No such data, however, are available for more recently developed subunit-selective GABA_A_R radioligands. To address this, we therefore combined a rat model of clinically relevant antipsychotic drug exposure with quantitative receptor autoradiography. Haloperidol (0.5 and 2 mg/kg/day) or drug vehicle were administered continuously to adult male Sprague-Dawley rats via osmotic mini-pumps for 28 days. Quantitative receptor autoradiography was then performed *post-mortem* using the GABA_A_R subunit-selective radioligand [^3^H]-Ro15-4513 and the non-subunit selective radioligand [^3^H]-flumazenil. Chronic haloperidol exposure increased [^3^H]-Ro15-4513 binding in the CA1 sub-field of the rat dorsal hippocampus (p<0.01; q<0.01; *d* = +1.3), which was not dose-dependent. [^3^H]-flumazenil binding also increased in most rat brain regions (p<0.05; main effect of treatment), irrespective of the haloperidol dose. These data confirm previous findings that chronic haloperidol exposure influences the specific binding of non-subtype selective GABA_A_R radioligands and is the first to demonstrate a potential effect of haloperidol on the binding of a α1/5GABA_A_R-selective radioligand. Although caution should be exerted when extrapolating results from animals to patients, our data support a view that exposure to antipsychotics may be a confounding factor in PET studies of GABA_A_R in the context of schizophrenia.

## Introduction

γ-Amino-butyric acid (GABA) is the major inhibitory neurotransmitter in the central nervous system (CNS). The GABA_A_ receptor (GABA_A_R) is a pentameric GABA-gated chloride ion channel composed of several classes of subunits (α1–6, β1–3, γ1–3, δ, θ, ρ, and □) (McKernan and Whiting, 1996). Of these, the diversity of the α-subunit is thought to be responsible for shaping the functional properties and ligand selectivity of the GABA_A_ benzodiazepine binding site (GABA_A_-BZR) (Barnard et al., 1998; Low et al., 2000; Mehta and Ticku, 1999; Tobler et al., 2001). Benzodiazepines act at the α/γ interface for the α subunits 1-3; 5 (Sigel and Steinmann, 2012). In the CNS, GABA_A_-BZR modulate pyramidal cell activity via tonic and phasic inhibition (Brickley and Mody, 2012; Mann and Paulsen, 2007; Whittington et al., 1995).

Deficits in GABA neurotransmission, resulting in disruptions to normal patterns of neural oscillatory activity are implicated in the pathophysiology of schizophrenia (Benes, 2010; Benes and Berretta, 2001; Lewis et al., 2012, 2005). In support of this, quantitative receptor autoradiography studies using [^3^H]-muscimol, an orthosteric agonist at the GABA binding site on GABA_A_-BZR, provide consistent evidence for increased binding density in frontal and temporal cortices and the caudate nucleus in *post-mortem* brain tissue from patients with schizophrenia (Benes et al., 1996; Dean et al., 1999; Deng and Huang, 2006; Hanada et al., 1987; Newell et al., 2007; Verdurand et al., 2013). By contrast, *post-mortem* studies focusing specifically on mRNA expression of GABA_A_ α-subunits report decreased expression of α1 (Beneyto et al., 2011; Glausier and Lewis, 2011), increased expression of α2 (Beneyto et al., 2011; Volk et al., 2002) and inconsistent results for the α5-subunit (Akbarian et al., 1995; Beneyto et al., 2011; Impagnatiello et al., 1998). A systematic review of positron emission tomography (PET) studies in schizophrenia patients using selective radiotracers for the BZ-site of the GABA_A_-BZR however found no consistent evidence for altered GABA_A_-BZR availability in schizophrenia (Egerton et al., 2017). Of note, these *post-mortem* data come from patients with a long duration of illness and exposure to antipsychotic medication. Similarly in most of the PET studies, the patients were also receiving antipsychotic medication (Egerton et al., 2017). Notably, different antipsychotics can directly alter the binding of ligands to GABA_A_-BZR, presumably by altering the expression and availability of the receptors (Frankle et al., 2015; Lee et al., 2013). Hence, distinguishing effect(s) of illness from antipsychotic exposure is challenging and medication may represent a significant source of heterogeneity in these data.

Rodent models offer the means to study direct effects of antipsychotic drugs on GABA_A_R radioligand binding, in the absence of illness effects and other confounds, and with strict control of genetic and environmental factors. Combining such models with quantitative receptor autoradiography (QAR) to assess drug influences on radioligand binding is also highly advantageous. Specifically, this method provides greater spatial resolution as compared to *in vivo* microPET, whilst keeping within a translational framework for comparison to clinical research using the same radioligands, which other techniques such as histology do not allow (Onwordi et al., 2020). Previous QAR studies in naïve adult rats provide evidence that chronic exposure to different antipsychotics influences the binding of both [^3^H]-muscimol (indexing GABA_A_R binding) and [^3^H]-flunitrazepam (indexing BZ-site binding) in a region-specific manner that is also dependent on the duration of drug exposure, the mode of administration and the sex of the animal (Dean et al., 2001; McLeod et al., 2008; See et al., 1990, 1989; Shirakawa and Tamminga, 1994; Skilbeck et al., 2008, 2007; Zink et al., 2004). For example, a 21 day oral exposure to aripiprazole (1mg/kg), olanzapine (1mg/kg) or risperidone (03.mg/kg) increased [^3^H]-muscimol binding in the striatum and nucleus accumbens of adolescent male rats, whilst in adolescent female rats binding was only increased in the prefrontal cortex (Lian and Deng, 2019). By contrast, a 12-day exposure to haloperidol (1 mg/kg/d) via intraperitoneal injections was reported to decrease [^3^H]-flumazenil binding in several regions of the rat brain (McLeod et al., 2008). The binding sites for the GABA_A_-BZR allosteric ligand, [^3^H]-flumazenil, in the rodent and human brain comprise both “zolpidem-sensitive” and “zolpidem-insensitive” sites, with the latter suggested to correspond to GABA_A_Rs that contain the α5 subunit (Mcleod et al., 2002). Consistent with the aforementioned data on [^3^H]-flumazenil binding, chronic (12d) systemic exposure to haloperidol resulted in a significant reduction in zolpidem-sensitive binding sites (α1,2,3GABA_A_R (Sancar et al., 2007), but had no effect on the insensitive-binding sites, suggesting a lack of effect of haloperidol on α5GABA_A_Rs (McLeod et al., 2008).

Notably, Kapur et al. reported that daily systemic (intraperitoneal) injections of antipsychotics, including haloperidol, result in no or inappropriately low occupancy at central dopamine D2 receptors at trough plasma levels (24 hours post-injection), since the half-life of antipsychotics in rodents is 4 to 6 times shorter as compared to humans (Kapur et al., 2003). This raises questions about the inferences drawn from these previous studies that have used doses unrepresentative of the clinical situation (Kapur et al., 2003). Furthermore, to date, no studies have examined the potential impact of antipsychotic drug exposure using radioligands with greater selectivity for GABA_A_-BZR containing α1/α5 subunits, such as Ro15-4513 (Lingford-hughes et al., 2002; Maeda et al., 2003). This is of clinical relevance, as a recent PET study in schizophrenia found reduced volume of distribution (V_T_) of [^11^C]-Ro15-4513 relative to healthy controls only in antipsychotic-naïve schizophrenia patients, whilst no group differences were found in antipsychotic-medicated schizophrenia patients relative to healthy controls (Marques et al., 2020). Furthermore, convergent lines of evidence from animal models strongly suggest that allosteric modulators of α5GABA_A_R have potential as novel, non-dopaminergic antipsychotic compounds, by balancing hippocampal excitation via tonic inhibition of pyramidal neurons (Bonin et al., 2007; Caraiscos et al., 2004; Donegan et al., 2019; Gerdjikov et al., 2008; Gill et al., 2011; Hauser et al., 2005; Semyanov et al., 2004; Towers et al., 2004). Importantly, the rescue of amphetamine-induced hyperlocomotion in the methylazoxymethanol acetate (MAM) neurodevelopmental disruption model of schizophrenia by the *α*5GABA_A_R positive allosteric modulator (PAM) SH-053-2’F-R-CH3, was abolished following prior exposure to the D2R antagonist haloperidol (Gill et al., 2014, 2011).

In the present study we therefore determined the impact of chronic exposure to haloperidol on GABA_A_R binding using *post-mortem* quantitative receptor autoradiography with [^3^H]-Ro15-4513 to assess *α*1/*α*5GABA_A_R and [^3^H]-flumazenil to assess BZ-sensitive *α*1-3;5GABA_A_R using a validated rat model of clinically comparable drug exposure (Kapur et al., 2003; Vernon et al., 2011). Based on the results of McLeod and colleagues (2008) who observed decreases in zolpidem-sensitive binding sites and no change in zolpidem-insensitive sites after haloperidol exposure, we hypothesized that chronic haloperidol exposure would decrease [^3^H]-flumazenil binding, with no effect on [^3^H]-Ro15-4513 binding.

## Experimental procedures

### Animals and treatment protocol

Male Sprague-Dawley rats (N=36, Charles River, UK; ∼ 10 weeks of age) were administered haloperidol (0.5 or 2 mg/kg/day; haloperidol; n=12/group: Sigma-Aldrich, Gillingham, Dorset, UK) or vehicle (β-hydroxypropylcyclodextrin, 20% w/v, acidified to pH 6 using ascorbic acid; n=12/group) using osmotic minipumps for 28 days (Vernon et al., 2011). Dyskinetic behavior, i.e., vacuous chewing movements, was assessed once at 26 days after the start of haloperidol exposure. This involved a simple measurement of purposeless chewing jaw movements in a 2-minute period, outside the home cage as described previously (Vernon et al., 2011). All experimental procedures were performed in accordance with the relevant guidelines and regulations, specifically, the Home Office (Scientific Procedures) Act 1986, United Kingdom and European Union (EU) directive 2010/63/EU and the approval of the local Animal Welfare and Ethical Review Body (AWERB) panels at King’s College London (for full details, see supplementary material).

### Quantitative receptor autoradiography with [^3^H]Ro15-4513 and [^3^H]flumazenil

On completion of drug or vehicle exposure, rats were culled by rising CO_2_ concentration and perfused transcardially with 100-200 ml of cold (+4°C) heparinized saline (50 IU/ml). The brains were then quickly dissected from the skull on a chilled platform, hemisected along the midline and flash frozen in isopentane. A plasma sample was collected from trunk blood for estimation of drug levels (see supplementary material). From one frozen hemisphere, coronal sections (20 µm-thick) from the left-brain hemisphere (Bregma +2.7 to −7.0mm) were cut in series using a cryostat (Leica CM1950), mounted onto glass slides (Superfrost™) and stored at −80°C until used for autoradiography. The remaining frozen hemisphere was utilized in separate experiments not related to this study.

[^3^H]-Ro15-4513 (Perkin Elmer, NET925250UC) was used to quantify α1/5GABA_A_R density. While this ligand has a high specificity (60-70%) (Myers et al., 2017) to α5GABA_A_R, a smaller proportion of the binding has affinity to α1GABA_A_R (Myers et al., 2012). Non-specific binding was determined by bretazenil (Sigma, B6434-25MG) due to its affinity to bind to a variety of GABA_A_R subtypes (α1-3;5) (Sieghart, 1995). Sections were pre-incubated at room temperature in Tris buffer (50 mM) for 20 minutes followed by incubation in either 2 nM [^3^H]-Ro15-4513 for specific binding, or 2 nM of [^3^H]-Ro15-4513 and 10 μM of bretazenil for non-specific binding at room temperature for 60 minutes. Slides were then washed in Tris buffer (2 × 2 min) at room temperature, dipped in distilled water (dH_2_0) and left to dry overnight. Dry slides were placed into light-tight cassettes with a radioactive [^3^H] standards slide, (ART-123A American Radiolabelled Chemicals Inc., USA) and hyperfilm (Amersham 8×10 in Hyperfilm Scientific Laboratory Supplies, UK). Films were exposed for 8 weeks before being developed in a Protex Ecomax film developer (Protec GmbH & Co, Germany). Identical procedures were used for [^3^H]-flumazenil (Perkin Elmer, NET757001MC), with the exception that the slides were incubated and washed in buffer at 4°C with 1nM [^3^H]-flumazenil, and 10 μM flunitrazepam (Sigma Aldrich, F-907 1ML) and 1nM [^3^H]-flumazenil, for specific and non-specific binding, respectively, and exposed for 4 weeks before development.

### Quantification of receptor binding

Films were developed and images were manually captured using a Nikon SLR camera and preprocessed (see supplementary material). The non-specific binding was negligible for both radioligands as illustrated in Figure S1. Optical density (OD) measurements were obtained using MCID software (Imaging Research Inc., 2003) from *a priori* defined regions of interest (ROIs; Fig. 1). These were chosen based on the known distribution of α1/α5GABA_A_R in the rat brain, data from prior studies reporting an effect of haloperidol on [^3^H]-muscimol or [^3^H]-flunitrazepam binding (McLeod et al., 2008; Skilbeck et al., 2008, 2007; Zink et al., 2004), and a defined role in the pathophysiology of schizophrenia (de la Fuente-Sandoval et al., 2017; Du and Grace, 2016; Gill et al., 2011; Heckers and Konradi, 2015; Lieberman et al., 2018; Miyamoto et al., 2005; Zink et al., 2004). Specific binding in nCi/mg was quantified using standard curves constructed from OD measurements of standards for each film, using the robust linear regression interpolation method in GraphPad (version 8.00, GraphPad Software, La Jolla California USA www.graphpad.com).

**Figure 1.**
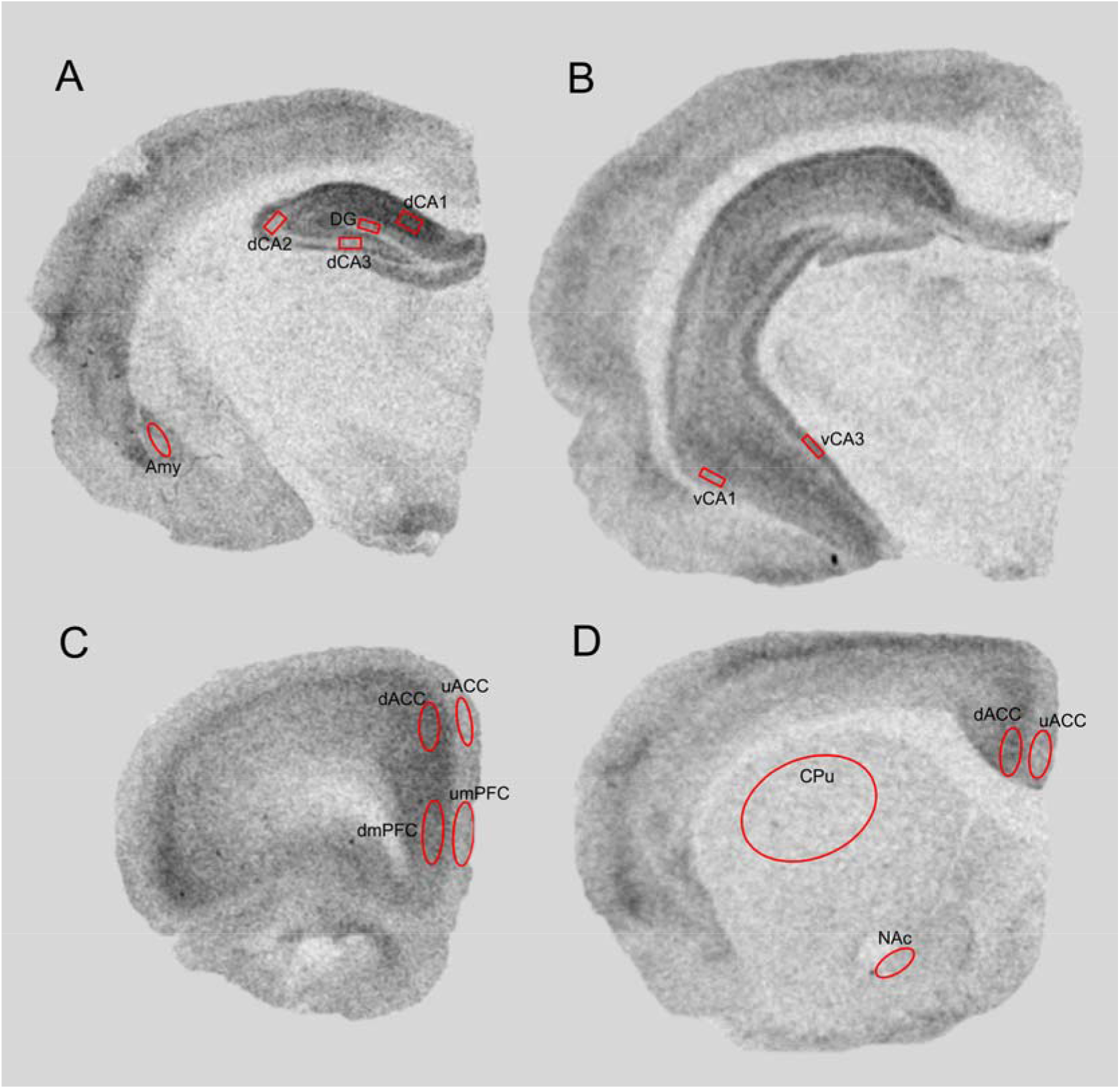
Representative [^3^H]-Ro15-4513 autoradiographs showing the placement of ROIs that were analyzed in this study. The same ROIs were used for the analysis of [^3^H]-flumazenil. A) dorsal hippocampal layers CA1 (dCA1), CA2 (dCA2), CA3 (dCA3), dentate gyrus (DG) and amygdala (amy). B) ventral hippocampal layers CA1 (vCA1), CA3 (vCA3). C) Medial Prefrontal Cortex (upper (1-3) and deeper (4-6) mPFC), Anterior Cingulate Cortex (upper (1-3) and deeper (4-6) ACC), differentiation of upper (1-3) vs deeper (4-6) layers was due to differential density of receptors across layers, specifically with α5 being more predominantly present in layer V and VI (Dunn et al., 1996). D) Caudate-Putamen (CPu), Nucleus Accumbens (NAc).

### Statistical Analyses

All statistical analyses were performed in Prism software (v8.0.0 for Macintosh, GraphPad Software, La Jolla California USA, www.graphpad.com). Tables 1 and 2 contain the final n-values per treatment group for each radioligand. In total, four rats were excluded from the complete dataset (n=2 for flumazenil and n=2 for Ro15-4513, respectively) for technical reasons. Specifically, data for one rat was missing due to a broken slide and artifacts were present in the films from the other three rats, which prevented the acquisition of images from these films for analysis. To check for the presence of possible outliers, we applied Grubb’s test once (not iteratively) to each dataset. Although some rats were identified as potential outliers from this analysis, we had no biological or technical reason to exclude them and they were thus included in the statistical analysis as described. The data were also checked for Gaussian distribution using the Shapiro-Wilk normality test. Group-level differences in ligand binding were assessed using a mixed-effects model, with ROI as within-subject factor and treatment (vehicle, haloperidol 0.5 or 2 mg/kg/day) as between-subject factor, using the specific binding (nCi/mg) of either [^3^H]-Ro15-4513 or [^3^H]-flumazenil as the dependent variable. Vacuous chewing movements scores were analyzed using non-parametric Kruskal-Wallis test (*p*<0.001). *Post-hoc* tests were performed where appropriate and corrected for multiple comparisons using the 2-stage set-up method of Benjamini, Krieger and Yekutieli, with the false discovery rate set at 5% (q<0.05) (Verhoeven et al., 2005). Relationships between vacuous chewing movements and ligand binding were modeled using non-parametric Spearman’s Rho correlation (2-tailed).

**Table 1.** Regional binding (nCi/mg) of [^3^H]-Ro15-4513 across the ROIs explored. Data shown are mean (SD, N), with p- and q-values (where q = FDR-adjusted p-value) for each ROI derived from *post-hoc* testing. Prefrontal Cortex (PFC), Anterior Cingulate Cortex (ACC), upper layer (1-3) and deeper layer (4-6); Caudate-Putamen (CPu), Nucleus Accumbens (NAc); dorsal hippocampal layers CA1 (dCA1), CA2 (dCA2), CA3 (dCA3), dentate gyrus (DG); ventral hippocampal layers CA1 (vCA1), CA3 (vCA3); amygdala (Amy). *Vehicle vs. Haloperidol 0.5 mg/kg/day (FDR q<0.01), # Haloperidol 0.5 mg/kg/day vs. Haloperidol 2 mg/kg/day (FDR q<0.01), ## (FDR q<0.001)

| ROI | Vehicle | HAL 0.5mg/kg/day | HAL 2mg/kg/day | Veh vs. HAL0.5mg/kg/day | Veh vs. HAL 2mg/kg/day | HAL0.5mg/kg/day vs. HAL2mg/kg/day |
| --- | --- | --- | --- | --- | --- | --- |
| Upper layer mPFC | 2.78 (1.28, 8) | 2.52 (1.06, 11) | 1.97 (1.26, 9) | $p=0.64$; $q=0.68$ | $p=0.21$; $q=0.50$ | $p=0.32$; $q=0.50$ |
| Deeper layer mPFC | 5.99 (2.06, 9) | 5.69 (1.38, 12) | 5.22 (1.34, 10) | $p=0.71$; $q=0.75$ | $p=0.35$; $q=0.66$ | $p=0.42$; $q=0.66$ |
| Upper layer ACC | 2.37 (1.02, 8) | 2.59 (1.19, 12) | 1.76 (1.08, 8) | $p=0.66$; $q=0.69$ | $p=0.27$; $q=0.42$ | $p=0.13$; $q=0.40$ |
| Deeper layer ACC | 4.85 (1.31, 9) | 4.66 (1.07, 12) | 4.61 (1.03, 10) | $p=0.72$; $q=0.96$ | $p=0.66$; $q=0.96$ | $p=0.92$; $q=0.96$ |
| CPU | 0.44 (0.34, 5) | 0.46 (0.40, 8) | 0.48 (0.39, 4) | $p=0.94$; $q=0.99$ | $p=0.90$; $q=0.99$ | $p=0.95$; $q=0.99$ |
| NAc | 1.54 (0.77, 7) | 2.30 (0.76, 9) | 0.80 (0.53, 7) ## | $p=0.07$; $q=0.049$ | $p=0.06$; $q=0.049$ | $p<0.001$; $q<0.001$ |
| dCA1 | 6.62 (1.05, 9) | 7.99 (1.09, 12) * | 6.66 (1.19, 10) # | $p<0.01$; $q<0.01$ | $p=0.94$; $q=0.33$ | $p<0.05$; $q<0.01$ |
| dCA2 | 4.45 (1.52, 9) | 4.56 (0.96, 12) | 3.72 (1.36, 10) | $p=0.86$; $q=0.90$ | $p=0.29$; $q=0.45$ | $p=0.12$; $q=0.38$ |
| DG | 3.26 (1.19, 9) | 3.44 (0.86, 12) | 3.13 (0.95, 10) | $p=0.71$; $q=0.84$ | $p=0.80$; $q=0.84$ | $p=0.45$; $q=0.84$ |
| dCA3 | 2.44 (0.83, 9) | 3.01 (0.85, 12) | 2.49 (0.86, 10) | $p=0.14$; $q=0.27$ | $p=0.90$; $q=0.94$ | $p=0.17$; $q=0.27$ |
| vCA1 | 4.85 (2.16, 3) | 3.71 (1.20, 12) | 2.94 (0.83, 9) | $p=0.46$; $q=0.48$ | $p=0.26$; $q=0.41$ | $p=0.10$; $q=0.31$ |
| vCA3 | 3.68 (3.12, 4) | 3.35 (1.65, 12) | 3.98 (1.02, 9) | $p=0.85$; $q=0.90$ | $p=0.86$; $q=0.90$ | $p=0.29$; $q=0.90$ |
| Amy | 2.11 (0.71, 9) | 1.99 (0.68, 12) | 1.83 (0.79, 10) | $p=0.72$; $q=0.76$ | $p=0.44$; $q=0.76$ | $p=0.61$; $q=0.76$ |

**Table 2.** Regional binding (nCi/mg) of [^3^H]-flumazenil across the ROIs explored. Data show mean (SD, N). Prefrontal Cortex (PFC), Anterior Cingulate Cortex (ACC), upper layer (1-3) and deeper layer (4-6); Caudate-Putamen (CPu), Nucleus Accumbens (NAc); dorsal hippocampal layers CA1 (dCA1), CA2 (dCA2),CA3 (dCA3), dentate gyrus (DG); ventral hippocampal layers CA1 (vCA1), CA3 (vCA3); amygdala (Amy).

| **[<sup>3</sup>H]Flumazenil binding** |  |  |  |
| --- | --- | --- | --- |
| **ROI** | **Vehicle** | **HAL 0.5mg/kg/day** | **HAL 2mg/kg/day** |
| **Upper layer mPFC** | **28.22 (8.2, 7)** | **35.43 (9.15, 11)** | **34.58 (7.95, 9)** |
| **Deeper layer mPFC** | **33.81 (7.91, 9)** | **40.62 (8.18, 11)** | **40.93 (8.37, 9)** |
| **Upper layer ACC** | **33.76 (5.70, 9)** | **39.04 (8.84, 12)** | **40.89 (8.39, 9)** |
| **Deeper layer ACC** | **27.05 (6.48, 9)** | **35.95 (7.52, 12)** | **35.55 (8.83, 9)** |
| **CPu** | **5.81 (2.83, 9)** | **8.62 (3.79, 12)** | **8.12 (4.56, 9)** |
| **NAc** | **10.75 (4.33, 9)** | **15.43 (5.83, 12)** | **18.99 (5.58, 9)** |
| **dCA1** | **29.30 (5.64, 9)** | **30.96 (7.40, 11)** | **34.54 (4.17, 10)** |
| **dCA2** | **15.89 (5.37, 9)** | **16.89 (5.27, 11)** | **19.54 (4.07, 10)** |
| **DG** | **26.57 (5.52, 9)** | **29.00 (5.84, 11)** | **32.58 (4.24, 10)** |
| **dCA3** | **17.25 (5.39, 9)** | **18.55 (5.36, 11)** | **22.10 (4.21, 10)** |
| **vCA1** | **14.81 (4.48, 5)** | **19.93 (7.07, 11)** | **21.81 (4.11, 9)** |
| **vCA3** | **14.24 (8.03, 5)** | **18.43 (7.53, 11)** | **24.24 (5.29, 9)** |
| **Amy** | **27.31 (8.11, 8)** | **31.08 (6.61, 11)** | **34.68 (8.48, 10)** |

## Results

### Haloperidol plasma levels and vacuous chewing movement behavior

Administration of haloperidol by osmotic pump achieved plasma levels (mean ± s.d.) of 2.96 ± 0.52 ng/mL and 12.2 ± 1.96 ng/mL, for the 0.5 and 2 mg/kg/day doses, respectively. Stereotypical vacuous chewing movement (VCM) behaviors were significantly different across treatment group (Kruskal-Wallis statistic = 9.98; p<0.001; Fig. S2). *Post-hoc* testing revealed a statistically significant increase in VCMs in rats exposed to 2 mg/kg/day haloperidol after 26 days exposure, relative to vehicle (p<0.01; q<0.05). There were no statistically significant differences between the haloperidol-exposed groups (p>0.05; q>0.05). The development of VCMs was not related to the binding of either [^3^H]-Ro15-4513 (Table S2) or [^3^H]-flumazenil across any ROI (Table S2). Haloperidol plasma levels were also significantly correlated with binding of either of the ligands used (Table S3).

### Effect of chronic haloperidol exposure on [^3^H]-Ro15-4513 specific binding measured with quantitative autoradiography

Mixed-effects model ANOVA revealed a statistically significant main effect of ROI (F(4,99)=105.0; p<0.0001) and ROI*treatment interaction (F(24,295)=1.70; p=0.02). No statistically significant main effect of treatment was observed (F(2,28)=1.28; p=0.30). *Post-hoc* testing on the ROI*treatment interaction revealed an increase in [^3^H]-Ro15-4513 specific binding only in the dCA1 of rats exposed to 0.5 mg/kg/day haloperidol, relative to both vehicle controls (p<0.01) and rats exposed to 2 mg/kg/d haloperidol (p<0.05) (Table 1; Figure 2 and 3A). These results remained significant after correction for multiple comparisons (FDR q<0.01) with a robust effect size (Cohen’s *d*= +1.3 and +1.2, respectively). There were no statistically significant differences between 2 mg/kg/day haloperidol exposed rats and vehicle controls (p=0.94; FDR q>0.05). In the NAc, exposure to 2 mg/kg/day haloperidol also decreased [^3^H]-Ro15-4513 binding relative to 0.5 mg/kg/day haloperidol-exposed rats (p<0.001; FDR q<0.001), but this failed to reach statistical significance with respect to vehicle controls (vehicle vs 0.5mg/kg/day: p=0.07; q=0.05; vehicle vs. 2mg/kg/day: p=0.06, q=0.05) (Table 1; Figure 2). All other ROIs showed no statistically significant changes in [^3^H]-Ro15-4513 binding after 28 days exposure to haloperidol, irrespective of the dose.

**Figure 2.**
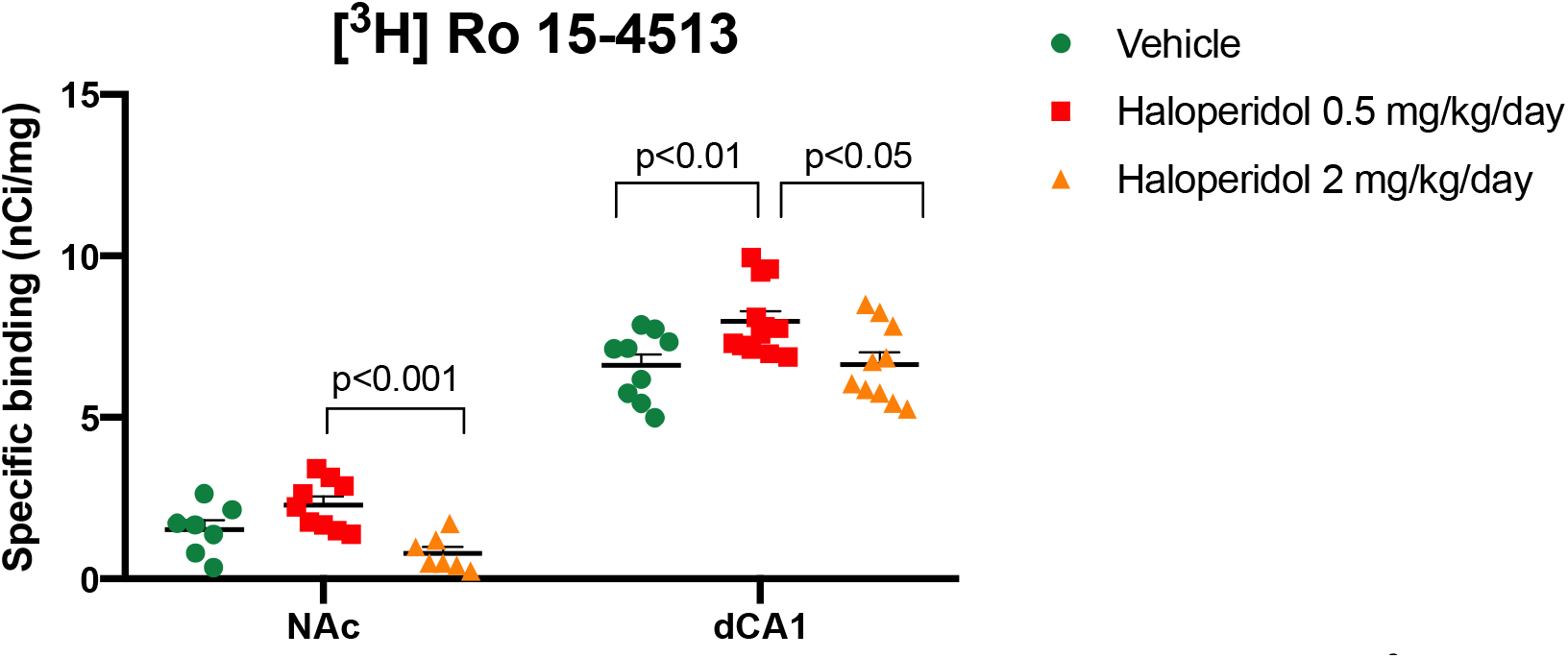
Chronic exposure to 0.5 mg/kg/d haloperidol results in elevated [^3^H]-Ro15-4513 specific binding in the rat nucleus accumbens (NAc) relative to the higher dose of haloperidol (2 mg/kg/d) and in the dorsal Cornu Ammonis 1 (dCA1) relative to both vehicle controls and the 2 mg/kg/d haloperidol group. Data points represent the specific binding values per individual animal (nCi/mg), horizontal line indicates group mean, and error bars indicate SEM. FDR-adjusted p-values are shown.

**Figure 3.**
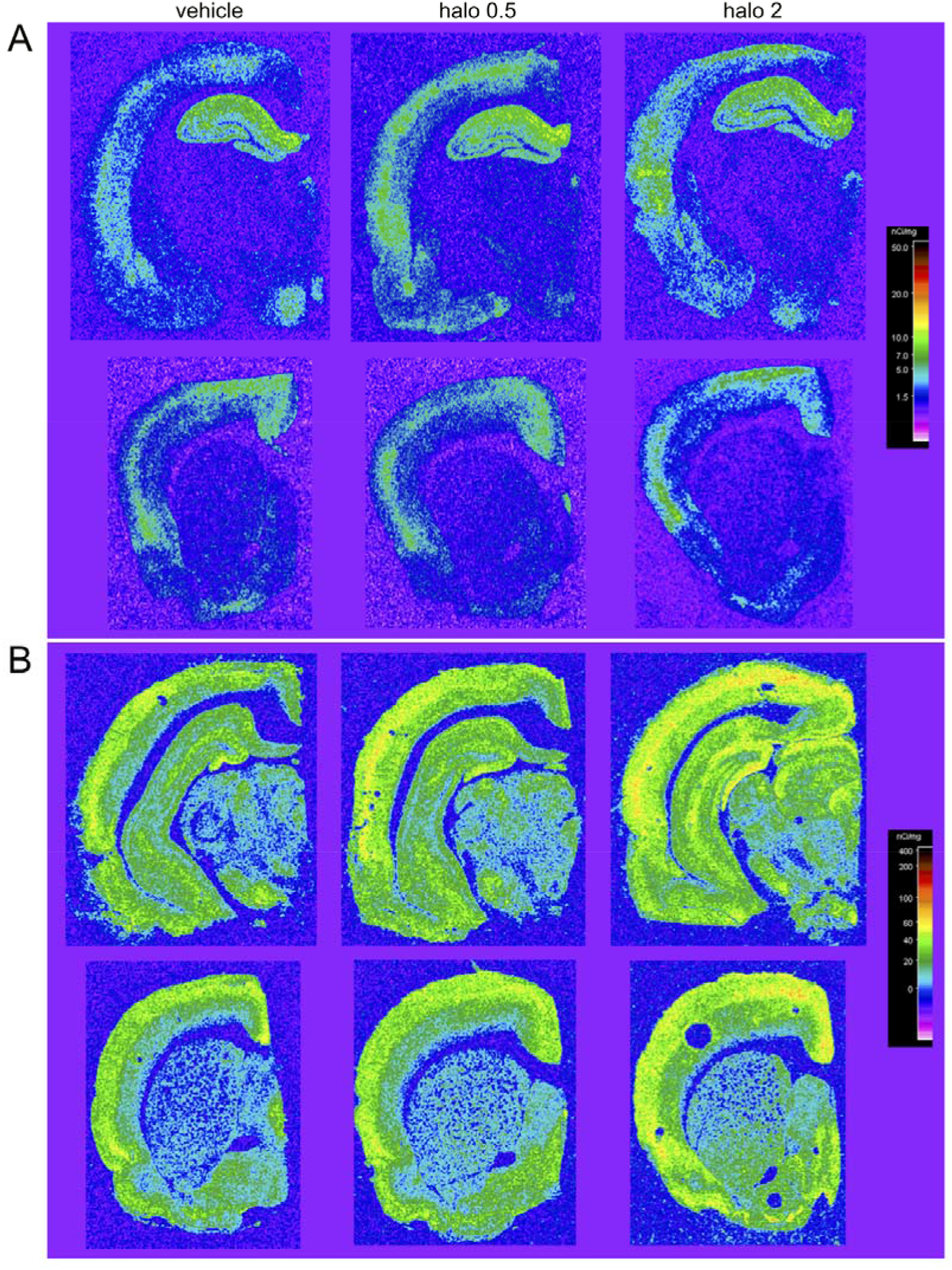
Pseudocolored representative autoradiographs showing A) [^3^H]-Ro15-4513 binding patterns and B) [^3^H]-flumazenil binding, by group: vehicle, haloperidol 0.5 mg/kg/day (halo 0.5), and haloperidol 2 mg/kg/day (halo 2). Non-specific binding for each ligand is shown in Figure S1.

### Effects of chronic haloperidol exposure on [^3^H]flumazenil binding measured with quantitative autoradiography

Mixed-effects model ANOVA revealed a significant main effect of ROI (F=(4.2, 107)=125; p<0.0001) (Table S4, Figure S3) and treatment (F(2, 28)=3.8; p=0.04), but no ROI*treatment interaction (F (24, 308)=1.1; p=0.3). Across all brain ROIs, the effect of haloperidol exposure was generally to increase [^3^H]-flumazenil-specific binding (see Table 2 and Figure 3B).

## Discussion

To our knowledge, this is the first study to investigate the effects of chronic (28d) exposure to haloperidol on GABA_A_R availability in a receptor subtype-specific manner using quantitative autoradiography. We observed that the specific binding of [^3^H]-Ro15-4513, a radioligand selective for α1/5-containing GABA_A_R is increased, but only in the dCA1 sub-region of the rat hippocampus an effect that was not dose-dependent.

By contrast, chronic exposure to haloperidol robustly increased the specific binding of the non-subtype selective GABA_A_R radioligand [^3^H]-flumazenil across the rat brain ROIs examined, irrespective of the dose administered. There were no statistically significant relationships between the specific binding of either radioligand in any brain ROI and either vacuous chewing movements, a proxy measure for haloperidol-induced tardive dyskinesia, or haloperidol plasma levels. Collectively, these data provide evidence that chronic exposure to haloperidol has a limited effect on the availability of α1/5-containing GABA_A_R, whilst it robustly affects the availability of α1-3;5 containing GABA_A_R as a whole in the naïve male rat brain.

Our findings were not consistent with our original hypotheses, based on prior work in this area. Specifically for [^3^H]-flumazenil, McLeod and Colleagues (2008) reported a decrease in the specific binding of this ligand in the rat frontal cortex, striatum, parietal cortex, dentate gyrus, CA1, 2 and 3 hippocampus subfields and thalamic nuclei following chronic haloperidol exposure (McLeod et al., 2008). By contrast, our data suggest a generalized increase in [^3^H]-flumazenil binding across these brain ROIs following chronic haloperidol exposure. Our data are however consistent with increased GABA_A_-BZR density in cortical areas measured using [^3^H]-flunitrazepam following chronic haloperidol exposure (Skilbeck et al., 2008, 2007). With regard to effects of haloperidol on α1/5-containing GABA_A_R, McLeod and colleagues (2008) reported no effect of this drug on zolpidem-insensitive [^3^H]-flumazenil binding, which is suggested to reflect α5GABA_A_R binding sites (Mcleod et al., 2002; McLeod et al., 2008). By contrast, we observed a localized effect of haloperidol on specific binding of the selective α1/5-containing GABA_A_R radioligand [^3^H]-Ro15-4513, only in the dCA1 of the rat hippocampus. Intriguingly, this was not dose-dependent and only occurred at the lower dose of haloperidol (0.5 mg/kg/d). Further experimentation is therefore required to both confirm this observation and understand its origin. The discrepancies between our findings and those of McLeod and colleagues (2008) are likely a reflection of methodological differences (McLeod et al., 2008). Specifically, our study utilized a selective α1/5GABA_A_R radioligand and the modes of antipsychotic dosing were different. Furthermore, there were variances between McLeod et al. (2008) and our work in the duration of drug exposure (12 vs. 28 days) and the age of the rats used (6 vs. 10 weeks of age). Consequently, further experimentation in younger and older animals are required with different durations and doses of haloperidol to unravel these complex effects of haloperidol GABA_A_R availability.

The mechanism driving drug-induced changes in GABA_A_R binding availability also remains unclear. Since haloperidol does not have affinity for the GABA_A_ receptor, it is likely that these are indirect as a result of the effects of haloperidol on neurotransmitter release through antagonism of dopamine D2-receptor (D2R) (McLeod et al., 2008). In this context, the role of dopamine in controlling GABA release, in which both D1 and D2R are involved, is well established (Starr, 1987). Increases in GABA also enhance the affinity of GABA_A_R for BZ-ligands such as flumazenil via a conformational change (Frankle et al., 2009; Miller et al., 1988; Tallman et al., 1978). In contrast, for BZR inverse agonists such as Ro15-4513, increased GABA levels appear to decrease the affinity of GABA_A_R for this ligand (Stokes et al., 2014). Changes in GABA release during haloperidol exposure thus likely account for the effect of haloperidol on GABA_A_R availability observed herein and by others (McLeod et al., 2008). Studies of bulk tissue GABA levels in the frontal cortex of schizophrenia patients using proton magnetic resonance spectroscopy (^1^H-MRS), however, report either no effect (Bojesen et al., 2019; Tayoshi et al., 2010) or a normalisation of elevated GABA levels (de la Fuente-Sandoval et al., 2017) following antipsychotic exposure, although this method only measures bulk tissue GABA and not synaptic levels. In contrast, rodent slice electrophysiology data suggest that D2R mediate synaptic GABA release onto pyramidal neurons in the PFC, whereby GABA release is decreased following dopamine administration (Xu and Yao, 2010). D2R antagonists, such as haloperidol, may therefore be predicted to increase GABA levels in the rodent frontal cortex, which could lead to elevated [^3^H]-flumazenil binding. In support of this view, GABA-immunoreactivity is increased in the axosomatic terminals of neurons in layers II, III, V, and VI in the frontal cortex of rats exposed chronically to 0.5 mg/kg/day haloperidol over 4 months (Vincent et al., 1994). On the other hand, an *in vivo* micro-dialysis study reported a decrease in the extracellular levels of GABA in the rat nucleus accumbens following chronic haloperidol exposure (See et al., 1992), which may perhaps suggest a compensatory upregulation of GABA_A_R in this region. Supporting these data, increased mRNA expression of GABA_A_R is reported in the rat NAc after chronic haloperidol exposure, (Pan et al., 2016b, 2016a). Further studies are however required using additional *in vivo* methods and *post-mortem* immunohistochemistry to understand how the effects of haloperidol on GABA_A_R availability observed herein relate to brain GABA and dopamine levels, and confirm these effects at the protein level, including localisation of these changes to specific cell types.

Some limitations of our study should be noted. First, while [^3^H]-Ro15-4513 binds predominantly to diazepam-sensitive GABA_A_R sites, it also binds to a diazepam-insensitive site in the cortex and hippocampus with lower affinity (Turner et al., 1991). This should be taken into consideration when comparing the binding patterns of [^3^H]-Ro15-4513 to those of the BZ-sensitive ligand [^3^H]-flumazenil. Second, we observed that in some brain regions, such as the corpus striatum, the [^3^H]-Ro15-4513 specific binding fell within the lower-range of the commercially available standards, limiting our ability to detect a specific signal from these regions with lower-binding values (likely reflecting lower receptor density). Hence, any effect of haloperidol may be underestimated or even missed in such regions. Third, we only examined the effects of haloperidol, a standard reference antipsychotic. Previous work suggests there may be differential effects of other antipsychotics with different receptor binding profiles, such as olanzapine and clozapine, on GABA_A_R availability in the rat brain (Farnbach-Pralong et al., 1998). In our previous studies comparing the effects of haloperidol and olanzapine on rat brain volume, radioligand binding (e.g. [^3^H]-UCB-J) and *post-mortem* cellular markers, we have consistently found no clear differences between these compounds (Cotel et al., 2015; Onwordi et al., 2020; Vernon et al., 2014). Whilst we therefore have no reason to believe that olanzapine for example, would not induce similar effects to haloperidol, this should be explicitly tested in future studies. Fourth, the effect of haloperidol on [^3^H]-Ro15-4513 binding in the dCA1 did not show true dose-dependence, with the effect only seen at the lower dose (0.5 mg/kg/d). The plasma levels of HAL in this group were slightly below the generally accepted therapeutic range (3 ng/ml c.f. 5-20 ng/ml), whilst the higher dose (2 mg/kg/d) fell within this, but had no effect on [^3^H]-Ro15-4513 binding in the dCA1. This study also had relatively small numbers and replication of this finding in a larger number of subjects will therefore be an important advance. Addressing the effect of sex as a biological variable is also necessary, since only male animals were used.

In conclusion, our findings suggest that exposure to haloperidol (and perhaps other antipsychotics) should be considered when measuring and interpreting global GABA_A_R binding availability using non-selective radioligands in the context of schizophrenia. Our data also provide initial evidence that haloperidol exposure has only a modest influence on α1/5-GABA_A_R selective radioligand binding. Of note, these data were collected in naïve animals, lacking any features relevant to the pathophysiology of schizophrenia. In this context, a recent PET study reported reduced [^11^C]-Ro15-4513 V_T_ in the hippocampus of antipsychotic-naïve schizophrenia patients, whilst no group differences were found in a second cohort of patients who were taking antipsychotics (Marques et al., 2020). One interpretation of these data is that antipsychotics may have a “normalizing” effect and it is tempting to speculate that this is consistent with the increases seen in GABA_A_R availability herein. Since the mechanism underlying these effects remains however, unknown, until this is better understood or convincingly refuted, one should be very cautious in drawing any clinical inferences from our data. Consequently, further studies to determine the effects of different antipsychotics on GABA_A_R availability in animal models reflective of genetic, environmental or pharmacological risk factors for schizophrenia are required to confirm or refute this hypothesis.

## Supporting information

Supl_Material_Peris-Yague et al 2020_revised

