## Supplementary material for "Region-specific and dose-specific effects of chronic haloperidol exposure on [^3^H]-Flumazenil and [^3^H]-Ro15-4513 GABA_A_ receptor binding sites in the rat brain": Supl_Material_Peris-Yague et al 2020_revised

United Kingdom

Dr Anthony C. Vernon

Department of Basic and Clinical Neuroscience,

Institute of Psychiatry, Psychology and Neuroscience,

King’s College London

Maurice Wohl Clinical Neuroscience Institute

5 Cutcombe Road

London SE5 9RT

United Kingdom

**Supplementary Methods**

*Animals*

All animals were housed in groups of four per cage in ventilated plastic cages (Tecniplast, UK) containing sawdust, paper sizzle nest and cardboard tunnels (Datesand group, UK) under a standard 12-hour light: dark cycle (lights on 07:00). The environment was maintained at 21±2°C, 55% ± 5% humidity. Animals had access to standard rat chow (Special Diet Services) and water *ad libitum* in the home cage. All procedures were done following the guidelines and regulations from the Home Office (Scientific Procedures) Act 1986, United Kingdom and European Union (EU) directive 2010/63/EU and the approval of the local Animal Welfare and Ethical Review Body (AWERB) panels at King’s College London.

### Rat model of clinically comparable antipsychotic drug dosing

The osmotic minipumps (AlzetModel 2ML4, 28 days; Alzet, Cupertino, California) filled with drug or vehicle solutions were inserted subcutaneously on the back flank under isoflurane anesthesia (5% induction, 1.5% maintenance). The doses of antipsychotic dosing were chosen based on previous D2R occupancy studies in our laboratory (Kapur et al., 2003); serum plasma levels achieved following chronic administration in this study reflect D_2_ occupancy in the range of 75% to 90% (Kapur et al., 2003; Vernon et al., 2011). The osmotic pump delivers at a steady rate in comparison with daily injections where drug levels fall to undetectable levels in 24 hours (half-life ~2.5 hours in rats for most antipsychotics).

*Quantification of haloperidol plasma levels*

Estimation of haloperidol plasma levels were obtained commercially using a contract research organization (Cyprotex, Macclesfield, UK). Briefly, plasma was collected by centrifugation of trunk blood samples obtained at the point of termination from each animal and 100 ul of each plasma sample shipped on dry ice along with 5 mg of haloperidol powder for standard curve calibration (Sigma-Aldrich, UK) to the CRO. The drug concentration in the plasma samples provided was then measured by LC-MS/MS, quantified against a calibration curve covering a dynamic range of 0.5 to 1000 ng/mL. The calibration standards, blanks, zeros (internal standard only), quality Control samples (3, 30 and 700 ng/mL) and rat plasma samples were mixed with organic solvent containing a mixture of three generic internal standards. Samples were treated at a ratio of 3 parts solvent to 1 part sample and, following protein precipitation, samples were either centrifuged (30 min, ca. 4 °C, 5000 g) or filtered through a 96‑well precipitation plate under positive pressure. The resulting supernatant/filtrate was then further diluted with HPLC grade water at a ratio of 2 parts HPLC grade water to 1 part supernatant/filtrate in a 96‑well plate. The plate was then sealed, vortex mixed and analyzed by LC-MS/MS with the amount of drug present in the plasma samples quantified from the calibration line and reported in ng/mL.

*Autoradiography development and image processing*

Developed films were placed on a light box (Northern lights, USA) and autoradiographs manually captured with a Nikon SLR camera and an AF-S Micro NIKKOR 60 mm F2.8G ED lens with lighting conditions kept constant. All images were converted to gray scale and the contrast normalized across each film by selecting the film background as white and the darkest standard as black using the ‘Levels’ tool and automated batch processing in Adobe Photoshop (Adobe Photoshop CC for Macintosh <https://www.adobe.com>).

**Supplementary Table 1.** Relationships between vacuous chewing movements and [^3^H]- Ro15-4513 binding across each region of interest. mPFC, medial prefrontal cortex; ACC, anterior cingulate cortex (upper layers (up), deeper layers (deep)); CPU, corpus striatum; NAc, nucleus accumbens; dCA, dorsal cornu ammonis 1,2,3; DG, dentate gyrus; vCA, ventral cornu ammonis 1,2,3; Amy, amygdala. *p<0.013 uncorrected for multiple comparisons, actual q-value = 0.21.

|  | **Group** | |  | |  | |
| --- | --- | --- | --- | --- | --- | --- |
|  | Vehicle | | Haloperidol (0.5 mg/kg/d) | | Haloperidol (2 mg/kg/d) | |
| **Region of interest** | Spearman’s ρ | *p*-value | Spearman’s ρ | *p*-value | Spearman’s ρ | *p*-value |
| up_mPFC | -.115 | .781 | .189 | .575 | -.059 | .879 |
| deep_mPFC | .364 | .327 | .124 | .700 | -.518 | .129 |
| up_ACC | .245 | .560 | .209 | .511 | .024 | .964 |
| deep_ACC | .174 | .652 | .166 | .603 | -.287 | .419 |
| CPU | .224 | .800 | -.491 | .223 | -.200 | .917 |
| Nac | .571 | .200 | .059 | .886 | .482 | .276 |
| dCA1 | .165 | .668 | -.191 | .548 | -.567 | .092 |
| dCA2 | -.028 | .956 | -.021 | .951 | -.439 | .204 |
| DG | .073 | .854 | -.708 | .013* | -.500 | .144 |
| dCA3 | -.073 | .854 | -.241 | .444 | -.274 | .440 |
| vCA1 | .866 | .667 | -.085 | .793 | -.277 | .465 |
| vCA3 | .775 | .500 | -.103 | .750 | -.017 | .974 |
| Amy | -.115 | .761 | -.089 | .784 | -.457 | .185 |

**Supplementary Table 2.** Relationships between vacuous chewing movements and [^3^H]-flumazenil binding across each region of interest. mPFC, medial prefrontal cortex; ACC, anterior cingulate cortex (upper layers (up), deeper layers (deep)); CPU, corpus striatum; NAc, nucleus accumbens; dCA, dorsal cornu ammonis 1,2,3; DG, dentate gyrus; vCA, ventral cornu ammonis 1,2,3; Amy, amygdala.

|  | **Group** | |  | |  | |
| --- | --- | --- | --- | --- | --- | --- |
|  | Vehicle | | Haloperidol (0.5 mg/kg/d) | | Haloperidol (2 mg/kg/d) | |
| **Region of interest** | Spearman’s ρ | *p*-value | Spearman’s ρ | *p*-value | Spearman’s ρ | *p*-value |
| up_mPFC | -.598 | .171 | .097 | .776 | .335 | .375 |
| deep_mPFC | -.516 | .156 | -.074 | .832 | -.243 | .528 |
| up_ACC | -.389 | .294 | -.170 | .595 | .326 | .390 |
| deep_ACC | -.507 | .167 | -.304 | .333 | -.184 | .634 |
| CPU | -.546 | .127 | -.032 | .925 | .142 | .717 |
| Nac | -.074 | .852 | -.173 | .587 | .519 | .156 |
| dCA1 | -.664 | .058 | .332 | .316 | .104 | .777 |
| dCA2 | -.304 | .423 | -.106 | .753 | .187 | .601 |
| DG | -.233 | .528 | -.203 | .547 | -.324 | .355 |
| dCA3 | -.175 | .643 | -.295 | .375 | -.409 | .241 |
| vCA1 | -.335 | .600 | -.433 | .183 | -.151 | .697 |
| vCA3 | -.894 | .100 | -.442 | .173 | .202 | .601 |
| Amy | -.206 | .643 | -.346 | .295 | -.445 | .198 |

**Supplementary Table 3.** Table showing the correlations between plasma levels achieved of haloperidol and binding of the ligands [^3^H]-Ro15-4513 and [^3^H]-flumazenil. Correlations were calculated using *Pearson’s Correlation* for normally distributed data except the following which were not normally distributed and thus we used *Spearman’s Correlation*: upper and deeper layers of the ACC and dCA1 in the [^3^H]Ro15-4513 haloperidol 0.5 mg/kg group; the NAc and dCA1 for the [^3^H]flumazenil haloperidol 0.5/mg/kg/day and the CPu for the 2 mg/kg/day dose of the same ligand.

|  | **[^3^H] Ro 15-4513** | | | | **[^3^H] Flumazenil** | | | |
| --- | --- | --- | --- | --- | --- | --- | --- | --- |
|  | Haloperidol (0.5 mg/kg/d) | | Haloperidol (2 mg/kg/d) | | Haloperidol (0.5 mg/kg/d) | | Haloperidol (2 mg/kg/d) | |
| **Region of interest** | Corr.coeficient | *p*-value | Corr.coeficient | *p*-value | Corr.coeficient | *p*-value | Corr. coeficient | *p*-value |
| up_mPFC | -0.23 | 0.5 | 0.61 | 0.08 | 0.06 | 0.86 | -0.08 | 0.84 |
| deep_mPFC | -0.49 | 0.11 | 0.48 | 0.16 | 0.25 | 0.47 | 0.41 | 0.27 |
| up_ACC | -0.070 | 0.835 | 0.43 | 0.29 | 0.26 | 0.42 | -0.28 | 0.47 |
| deep_ACC | -0.24 | 0.44 | 0.54 | 0.11 | 0.54 | 0.07 | 0.38 | 0.31 |
| CPU | 0.52 | 0.18 | -0.02 | 0.98 | 0.18 | 0.58 | -0.5 | 0.91 |
| NAc | -0.66 | 0.06 | -0.09 | 0.85 | 0.31 | 0.33 | -0.19 | 0.62 |
| dCA1 | -0.08 | 0.82 | 0.56 | 0.09 | 0.05 | 0.90 | -0.08 | 0.82 |
| dCA2 | 0.06 | 0.86 | 0.26 | 0.47 | 0.10 | 0.77 | -0.14 | 0.69 |
| DG | 0.22 | 0.48 | 0.66 | 0.04 | 0.24 | 0.47 | 0.27 | 0.44 |
| dCA3 | -0.25 | 0.43 | 0.31 | 0.38 | 0.23 | 0.50 | 0.14 | 0.69 |
| vCA1 | -0.24 | 0.45 | 0.46 | 0.21 | 0.15 | 0.66 | 0.12 | 0.75 |
| vCA3 | -0.66 | 0.02 | -0.24 | 0.54 | 0.22 | 0.51 | -0.17 | 0.67 |
| Amy | -0.25 | 0.44 | 0.39 | 0.27 | 0.06 | 0.86 | 0.23 | 0.52 |

**Supplementary Table 4.** Comparison of the means of the ROIs of [^3^H]-flumazenil. Results show differences in binding across ROIs. FDR post-hoc testing revealed differences across most ROIs (q<0.05) except: upper layer mPFC vs. deeper layer ACC (p=0.93, q>0.05), upper layer mPFC vs. dCA1 (p=0.33, q=0.5), deeper layer mPFC vs upper layer ACC (p=0.56, q>0.05), deeper layer ACC vs. dCA1 (p=0.33, q=0.05), dCA1 vs amyg (p=0.79, q>0.05), dCA3 vs vCA1 (p=0.75, q>0.05), dCA3 vs. vCA1 (p=0.75, q>0.05) and vCA1 vs vCA3 (p=0.92, q>0.05).

| **Test details** | **Mean 1** | **Mean 2** | **Mean Diff** | **SE of diff** | **N1** | **N2** | **t** | **DF** |
| --- | --- | --- | --- | --- | --- | --- | --- | --- |
| up_mPFC vs. down_mPFC | 33.28 | 38.61 | -5.328 | 0.8941 | 27 | 29 | 5.958 | 26.00 |
| up_mPFC vs. up_ACC | 33.28 | 38.01 | -4.737 | 0.9520 | 27 | 30 | 4.976 | 26.00 |
| up_mPFC vs. down_ACC | 33.28 | 33.16 | 0.1139 | 1.352 | 27 | 30 | 0.08428 | 26.00 |
| up_mPFC vs. CPU_str | 33.28 | 7.625 | 25.65 | 1.244 | 27 | 30 | 20.62 | 26.00 |
| up_mPFC vs. Nac_vStr | 33.28 | 15.09 | 18.19 | 1.235 | 27 | 30 | 14.73 | 26.00 |
| up_mPFC vs. dCA1 | 33.28 | 31.66 | 1.619 | 1.626 | 27 | 30 | 0.9955 | 25.00 |
| up_mPFC vs. dCA2 | 33.28 | 17.47 | 15.80 | 1.546 | 27 | 30 | 10.22 | 25.00 |
| up_mPFC vs. DG | 33.28 | 29.46 | 3.813 | 1.498 | 27 | 30 | 2.546 | 25.00 |
| up_mPFC vs. dCA3 | 33.28 | 19.34 | 13.93 | 1.495 | 27 | 30 | 9.322 | 25.00 |
| up_mPFC vs. vCA1 | 33.28 | 19.58 | 13.70 | 1.800 | 27 | 25 | 7.611 | 20.00 |
| up_mPFC vs. vCA3 | 33.28 | 19.69 | 13.59 | 1.791 | 27 | 25 | 7.588 | 20.00 |
| up_mPFC vs. Amyg | 33.28 | 31.28 | 1.996 | 1.881 | 27 | 29 | 1.061 | 24.00 |
| down_mPFC vs. up_ACC | 38.61 | 38.01 | 0.5908 | 1.004 | 29 | 30 | 0.5886 | 28.00 |
| down_mPFC vs. down_ACC | 38.61 | 33.16 | 5.442 | 0.7773 | 29 | 30 | 7.001 | 28.00 |
| down_mPFC vs. CPU_str | 38.61 | 7.625 | 30.98 | 1.122 | 29 | 30 | 27.61 | 28.00 |
| down_mPFC vs. Nac_vStr | 38.61 | 15.09 | 23.52 | 1.101 | 29 | 30 | 21.36 | 28.00 |
| down_mPFC vs. dCA1 | 38.61 | 31.66 | 6.946 | 1.485 | 29 | 30 | 4.678 | 27.00 |
| down_mPFC vs. dCA2 | 38.61 | 17.47 | 21.13 | 1.460 | 29 | 30 | 14.47 | 27.00 |
| down_mPFC vs. DG | 38.61 | 29.46 | 9.141 | 1.227 | 29 | 30 | 7.448 | 27.00 |
| down_mPFC vs. dCA3 | 38.61 | 19.34 | 19.26 | 1.323 | 29 | 30 | 14.56 | 27.00 |
| down_mPFC vs. vCA1 | 38.61 | 19.58 | 19.02 | 1.547 | 29 | 25 | 12.29 | 22.00 |
| down_mPFC vs. vCA3 | 38.61 | 19.69 | 18.92 | 1.568 | 29 | 25 | 12.07 | 22.00 |
| down_mPFC vs. Amyg | 38.61 | 31.28 | 7.324 | 1.753 | 29 | 29 | 4.178 | 26.00 |
| up_ACC vs. down_ACC | 38.01 | 33.16 | 4.851 | 1.243 | 30 | 30 | 3.902 | 29.00 |
| up_ACC vs. CPU_str | 38.01 | 7.625 | 30.39 | 1.037 | 30 | 30 | 29.32 | 29.00 |
| up_ACC vs. Nac_vStr | 38.01 | 15.09 | 22.92 | 0.9332 | 30 | 30 | 24.57 | 29.00 |
| up_ACC vs. dCA1 | 38.01 | 31.66 | 6.355 | 1.438 | 30 | 30 | 4.419 | 28.00 |
| up_ACC vs. dCA2 | 38.01 | 17.47 | 20.54 | 1.484 | 30 | 30 | 13.84 | 28.00 |
| up_ACC vs. DG | 38.01 | 29.46 | 8.550 | 1.310 | 30 | 30 | 6.526 | 28.00 |
| up_ACC vs. dCA3 | 38.01 | 19.34 | 18.67 | 1.393 | 30 | 30 | 13.40 | 28.00 |
| up_ACC vs. vCA1 | 38.01 | 19.58 | 18.43 | 1.631 | 30 | 25 | 11.30 | 23.00 |
| up_ACC vs. vCA3 | 38.01 | 19.69 | 18.33 | 1.594 | 30 | 25 | 11.50 | 23.00 |
| up_ACC vs. Amyg | 38.01 | 31.28 | 6.733 | 1.859 | 30 | 29 | 3.621 | 27.00 |
| down_ACC vs. CPU_str | 33.16 | 7.625 | 25.54 | 1.113 | 30 | 30 | 22.94 | 29.00 |
| down_ACC vs. Nac_vStr | 33.16 | 15.09 | 18.07 | 1.099 | 30 | 30 | 16.45 | 29.00 |
| down_ACC vs. dCA1 | 33.16 | 31.66 | 1.505 | 1.530 | 30 | 30 | 0.9833 | 28.00 |
| down_ACC vs. dCA2 | 33.16 | 17.47 | 15.69 | 1.471 | 30 | 30 | 10.67 | 28.00 |
| down_ACC vs. DG | 33.16 | 29.46 | 3.699 | 1.228 | 30 | 30 | 3.013 | 28.00 |
| down_ACC vs. dCA3 | 33.16 | 19.34 | 13.82 | 1.311 | 30 | 30 | 10.54 | 28.00 |
| down_ACC vs. vCA1 | 33.16 | 19.58 | 13.58 | 1.469 | 30 | 25 | 9.246 | 23.00 |
| down_ACC vs. vCA3 | 33.16 | 19.69 | 13.48 | 1.623 | 30 | 25 | 8.305 | 23.00 |
| down_ACC vs. Amyg | 33.16 | 31.28 | 1.882 | 1.776 | 30 | 29 | 1.060 | 27.00 |
| CPU_str vs. Nac_vStr | 7.625 | 15.09 | -7.464 | 0.6687 | 30 | 30 | 11.16 | 29.00 |
| CPU_str vs. dCA1 | 7.625 | 31.66 | -24.03 | 1.094 | 30 | 30 | 21.97 | 28.00 |
| CPU_str vs. dCA2 | 7.625 | 17.47 | -9.849 | 0.9473 | 30 | 30 | 10.40 | 28.00 |
| CPU_str vs. DG | 7.625 | 29.46 | -21.84 | 0.8894 | 30 | 30 | 24.55 | 28.00 |
| CPU_str vs. dCA3 | 7.625 | 19.34 | -11.72 | 0.8371 | 30 | 30 | 14.00 | 28.00 |
| CPU_str vs. vCA1 | 7.625 | 19.58 | -11.96 | 0.9780 | 30 | 25 | 12.22 | 23.00 |
| CPU_str vs. vCA3 | 7.625 | 19.69 | -12.06 | 1.293 | 30 | 25 | 9.329 | 23.00 |
| CPU_str vs. Amyg | 7.625 | 31.28 | -23.66 | 1.461 | 30 | 29 | 16.19 | 27.00 |
| Nac_vStr vs. dCA1 | 15.09 | 31.66 | -16.57 | 1.282 | 30 | 30 | 12.93 | 28.00 |
| Nac_vStr vs. dCA2 | 15.09 | 17.47 | -2.385 | 1.071 | 30 | 30 | 2.228 | 28.00 |
| Nac_vStr vs. DG | 15.09 | 29.46 | -14.37 | 0.9848 | 30 | 30 | 14.60 | 28.00 |
| Nac_vStr vs. dCA3 | 15.09 | 19.34 | -4.255 | 0.9133 | 30 | 30 | 4.659 | 28.00 |
| Nac_vStr vs. vCA1 | 15.09 | 19.58 | -4.491 | 0.9646 | 30 | 25 | 4.656 | 23.00 |
| Nac_vStr vs. vCA3 | 15.09 | 19.69 | -4.596 | 1.289 | 30 | 25 | 3.564 | 23.00 |
| Nac_vStr vs. Amyg | 15.09 | 31.28 | -16.19 | 1.484 | 30 | 29 | 10.91 | 27.00 |
| dCA1 vs. dCA2 | 31.66 | 17.47 | 14.18 | 0.7999 | 30 | 30 | 17.73 | 29.00 |
| dCA1 vs. DG | 31.66 | 29.46 | 2.194 | 0.7678 | 30 | 30 | 2.858 | 29.00 |
| dCA1 vs. dCA3 | 31.66 | 19.34 | 12.31 | 0.8560 | 30 | 30 | 14.39 | 29.00 |
| dCA1 vs. vCA1 | 31.66 | 19.58 | 12.08 | 1.164 | 30 | 25 | 10.37 | 24.00 |
| dCA1 vs. vCA3 | 31.66 | 19.69 | 11.97 | 0.8295 | 30 | 25 | 14.44 | 24.00 |
| dCA1 vs. Amyg | 31.66 | 31.28 | 0.3774 | 1.408 | 30 | 29 | 0.2681 | 28.00 |
| dCA2 vs. DG | 17.47 | 29.46 | -11.99 | 0.6596 | 30 | 30 | 18.18 | 29.00 |
| dCA2 vs. dCA3 | 17.47 | 19.34 | -1.870 | 0.4018 | 30 | 30 | 4.654 | 29.00 |
| dCA2 vs. vCA1 | 17.47 | 19.58 | -2.106 | 0.7909 | 30 | 25 | 2.663 | 24.00 |
| dCA2 vs. vCA3 | 17.47 | 19.69 | -2.211 | 0.9638 | 30 | 25 | 2.294 | 24.00 |
| dCA2 vs. Amyg | 17.47 | 31.28 | -13.81 | 1.137 | 30 | 29 | 12.14 | 28.00 |
| DG vs. dCA3 | 29.46 | 19.34 | 10.12 | 0.5834 | 30 | 30 | 17.35 | 29.00 |
| DG vs. vCA1 | 29.46 | 19.58 | 9.884 | 0.7688 | 30 | 25 | 12.86 | 24.00 |
| DG vs. vCA3 | 29.46 | 19.69 | 9.779 | 0.8945 | 30 | 25 | 10.93 | 24.00 |
| DG vs. Amyg | 29.46 | 31.28 | -1.817 | 1.112 | 30 | 29 | 1.634 | 28.00 |
| dCA3 vs. vCA1 | 19.34 | 19.58 | -0.2365 | 0.7219 | 30 | 25 | 0.3276 | 24.00 |
| dCA3 vs. vCA3 | 19.34 | 19.69 | -0.3408 | 0.9183 | 30 | 25 | 0.3712 | 24.00 |
| dCA3 vs. Amyg | 19.34 | 31.28 | -11.94 | 1.044 | 30 | 29 | 11.43 | 28.00 |
| vCA1 vs. vCA3 | 19.58 | 19.69 | -0.1044 | 0.9858 | 25 | 25 | 0.1059 | 24.00 |
| vCA1 vs. Amyg | 19.58 | 31.28 | -11.70 | 1.413 | 25 | 29 | 8.283 | 24.00 |
| vCA3 vs. Amyg | 19.69 | 31.28 | -11.60 | 1.469 | 25 | 29 | 7.896 | 24.00 |

**Supplementary Figure 1.** Representative images of autoradiographs from one animal processed for [^3^H]-Ro15-4513 and one for [^3^H]-Flumazenil. Specific binding (SB) is shown on the top and corresponding non-specific binding (NSB) on the bottom. Since NSB was completely absent, we did not measure it.

**Supplementary Figure 2.** Vacuous chewing movements score in each treatment group (vehicle vs. haloperidol 0.5 mg/kg/d vs. haloperidol 2mg/kg/d). Kruskal-Wallis statistic=9.98; p<0.001 revealed differences across all groups. *Post-hoc* test showed a significant increase in vacuous chewing movements in the haloperidol 2mg/kg/d group vs. vehicle group (p<0.01; q<0.05).

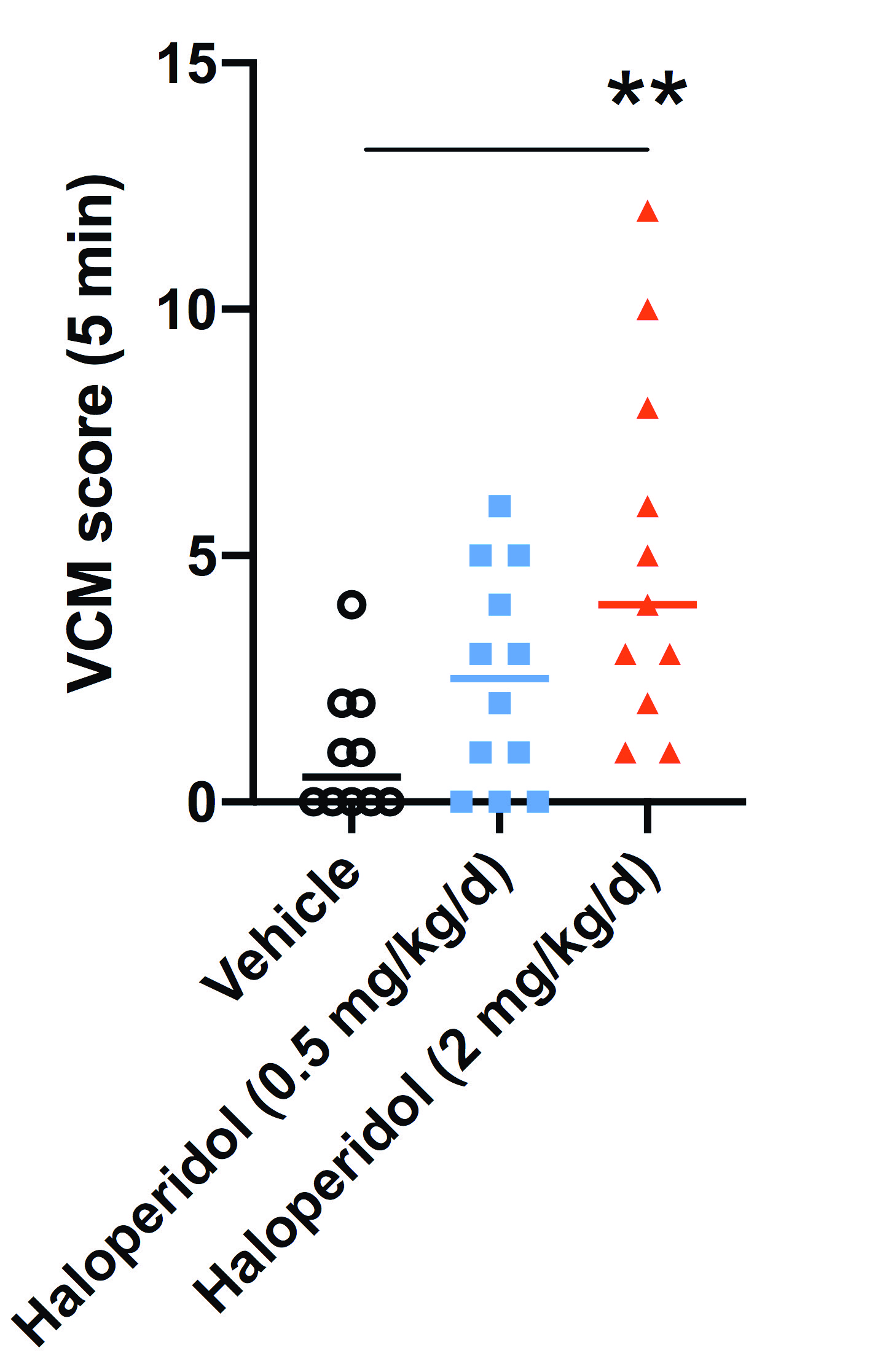

**Supplementary Figure 3.** [^3^H]flumazenil binding pattern in all the ROIs regardless of treatment. Prefrontal Cortex (PFC), Anterior Cingulate Cortex (ACC), upper layer (1-3) and deeper layer (4-6); Caudate-Putamen (CPu), Nucleus Accumbens (NAc); dorsal hippocampal layers CA1 (dCA1), CA2 (dCA2),CA3 (dCA3), dentate gyrus (DG); ventral hippocampal layers CA1 (vCA1), CA3 (vCA3), amygdala (Amy).

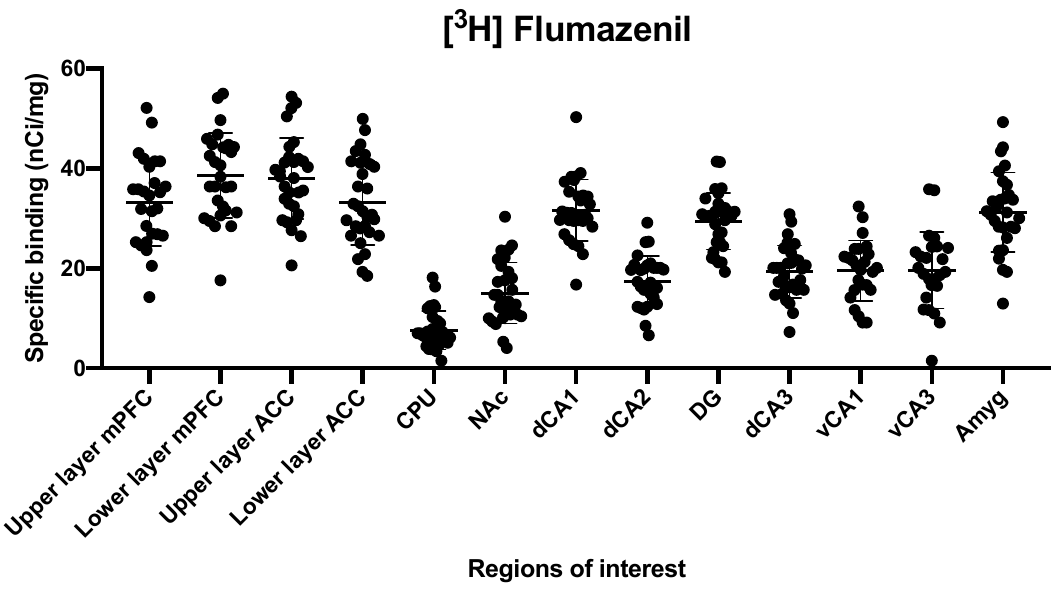
